# Hif-1α stabilisation polarises macrophages via cyclooxygenase/prostaglandin E2 *in vivo*

**DOI:** 10.1101/536862

**Authors:** Amy Lewis, Philip M. Elks

**Author notes:** Corresponding author: Dr Philip M. Elks The Bateson Centre, University of Sheffield, Firth Court, Western Bank, Sheffield, South Yorkshire, S10 2TN. UK.

## Abstract

Macrophage subtypes are poorly characterised in disease systems *in vivo*. The initial innate immune response to injury and infectious stimuli through M1 polarisation is important for the outcome of disease. Appropriate macrophage polarisation requires complex coordination of local microenvironmental cues and cytokine signalling to influence immune cell phenotypes. If the molecular mechanisms behind macrophage polarisation were better understood then macrophages could be pharmacologically tuned to better deal with bacterial infections, for example tuberculosis. Here, using zebrafish *tnfa:GFP* transgenic lines as *in vivo* readouts of M1 macrophages, we show that hypoxia and stabilisation of Hif-1α polarises macrophages to a *tnfa* expressing phenotype. We demonstrate a novel mechanism of Hif-1α mediated macrophage *tnfa* upregulation via a cyclooxygenase/prostaglandin E2 axis, a mechanism that is conserved in human primary macrophages. These findings uncover a novel macrophage HIF/COX/TNF axis that links microenvironmental cues to macrophage phenotype that may have implications in inflammation, infection and cancer, where hypoxia is a common microenvironmental feature and where cyclooxygenase and Tnfa are major mechanistic players.

## Introduction

The innate immune response to injury and pathogen invasion are complex and are tightly regulated by coordination of microenvironmental cues and cytokine signalling. Macrophages are important innate immune cells in disease and their activation states, commonly termed polarisation states, are especially important during the early response to injury and infection. Mammalian macrophages have been classified into two broad polarisation/activation states; M1 (or classically activated) and M2 (alternatively activated) (1). Pro-inflammatory, or M1, macrophages are highly antimicrobial and can phagocytose and efficiently kill bacteria. Macrophages are also central players in tissue healing and restoration of homeostasis post injury/infection, which requires a change in macrophage phenotype to a wound healing, M2, state (2). A diverse variety in macrophage function has been observed in many *in vitro* systems, yet control of macrophage subsets remains poorly understood *in vivo*. Current classifications are based on *in vitro* mammalian datasets, using monocytes that have been polarised to different macrophage phenotypes by addition of cytokine-cocktails to the media (3). Specific subsets of macrophage polarisation are context dependent *in vivo* and subtle changes in the microenvironment can lead to different phenotypes of M1 and M2 macrophages, reflecting a spectrum rather than a binary classification of behaviours (4). One important microenvironmental factor is hypoxia, which has profound effects on macrophage phenotypes (5). To fully understand how microenvironmental cues affect macrophage polarisation, *in vivo* systems are required to study macrophage behaviour over time and space without disturbance from extrinsic factors. Determining the molecular triggers and polarisation states of macrophages could open up possibilities to manipulate macrophage phenotype during disease as novel therapeutic avenues.

The tissue macrophage response to injury/bacterial infections is a rapid, pathogen recognition receptor (PPR) mediated, switch to an M1 type phenotype, characterised by production and release of pro-inflammatory mediators, including cytokines such as IL-1β, IL-6, IL-12 and TNF (6, 7). This phenotypic switch is required for a successful response to the invading pathogen and for clearance before infection can take hold (8, 9). However, in the case of intracellular pathogens, such as *Mycobacterium tuberculosis* (Mtb), this macrophage response is subverted, establishing a protective niche within innate immune cells (10–13). This bacterial-mediated interference in macrophage polarisation highlights a gap for potential therapeutic tuning of the macrophage response to allow for bacterial clearance (14, 15).

One potential macrophage tuning mechanism is via the Hypoxia Inducible Factor (HIF) −1α transcription factor (16). HIF-1α is a master transcriptional regulator of the cellular response to low levels of oxygen (hypoxia) (17). Leukocytes, including macrophages and neutrophils, are exquisitely sensitive to tissue hypoxia, a signature of many disease microenvironments (eg infections and cancer) due to lack of local blood supply and a high turnover of local oxygen by pathogens (5, 18). Hypoxia and HIF-1α stabilisation have been shown to have activating effects on macrophages *in vitro*, increasing their M1 profile and bactericidal capabilities, yet the pro-inflammatory mechanisms behind this are not well understood *in vivo* (5).

Hypoxia is a microenvironmental signal that increases epithelial cyclooxygenase/prostaglandin E2 production in cancer models (19–21). Eicosanoids are lipid signalling molecules and important pro-inflammatory mediators which are produced, and released, by macrophages during early microbial infection (22, 23). The best characterised of these are prostaglandins, strong inflammatory mediators that are known to affect macrophage polarisation states during infection (24). A key family of enzymes for prostaglandin biosynthesis is the cyclooxygenases (COX). COX-1 and -2 enzymes are responsible for the production of prostaglandins as breakdown products of arachidonic acid.

Prostaglandins, in turn, are highly immunomodulatory, synergising with cytokines to amplify pro-inflammatory responses (25). The potential effects of hypoxia-induced prostaglandins on macrophage polarisation have not been investigated *in vivo*. Recent evidence indicates that a HIF/COX/TNF axis may exist, with hypoxia-induced TNF expression in osteoblasts (bone producing cells) being mediated via cyclooxygenase enzymes by an, as yet, unknown mechanism (26). If observed in macrophages, this HIF/COX/TNF axis could have important pro-inflammatory outcomes in disease and represent a novel therapeutic avenue.

Zebrafish have emerged as a useful model to understand innate immune cells in their *in vivo* microenvironment (27). Development of multiple innate immune cell transgenic lines, combined with non-invasive fluorescence imaging, has enabled mechanistic investigation into macrophage biology (2, 28, 29). Well-characterised models of inflammation and infection biology have emerged using zebrafish larvae. Mechanistic insights into neutrophil and macrophage recruitment to an injury site have been gained through use of a tailfin transection model (30), while host-pathogen interactions have been studied using the natural fish pathogen *Mycobacterium marinum* (Mm) as a model of tuberculosis (27). Recently, it has been demonstrated that *tnfa* promoter driven fluorescent transgenic lines display an M1 type phenotype upon sterile injury immune challenge (31, 32). This advance has enabled us to investigate macrophage polarisation within the zebrafish system during a host-pathogen immune challenge. We have previously demonstrated that Hif-1α stabilisation induces macrophage proinflammatory *il-1β* transcription and that this is host-protective against Mm infection in the zebrafish model (33). We therefore hypothesised that Hif-1αstabilisation would promote macrophage pro-inflammatory *tnfa* expression, via a cyclooxygenase dependent mechanism, that could also be beneficial to the host during infection.

Here, we investigated the regulation of expression of the important M1 pro-inflammatory cytokine *tnfa* by Hif-1α stabilisation *in vivo*. We demonstrate that genetic, pharmacological and hypoxia-mediated Hif-1α stabilisation upregulated macrophage *tnfa* in zebrafish. We show that *tnfa* transcription is upregulated in macrophages after injury and Mm infection (both at early- and granuloma- stage infection). We identify that Hif-1α activates macrophage *tnfa* via cyclooxygenase/prostaglandin E2, while injury/Mm driven *tnfa* production does not require active cyclooxygenase. Importantly, this novel macrophage HIF/COX/TNF axis is conserved in primary human macrophages. These findings have important implications in inflammation, infection and cancer biology where macrophage polarisation states are influenced by microenvironmental hypoxia.

## Materials and Methods

### Zebrafish

Zebrafish were raised and maintained on a 14:10-hour light/dark cycle at 28°C, according to standard protocols (34), in UK Home Office approved facilities at The Bateson Centre aquaria at the University of Sheffield. Strains used were Nacre (wildtype), *TgBAC(tnfa:GFP)pd1028*, *Tg(tnfa:eGFP-F)ump5Tg*, *Tg(mpeg1:mCherry-F)ump2Tg, Tg(mpeg1:mCherryCAAX)sh378, Tg(lyz:Ds-RED2)nz50 TgBAC(il-1β:eGFP)sh445* and *Tg(phd3:EGFP)i144* (31, 32, 35–37).

### Tailfin injury

Inflammation was induced in zebrafish embryos by tail transection as previously described (38). Embryos were anaesthetised at 2 days post fertilisation (dpf) by immersion in 0.168mg/ml Tricaine, and transection of the tail was performed using a scalpel blade. Larvae were imaged by confocal microscopy at 16 hour post wound (hpw) on a Leica TCS-SPE confocal on an inverted Leica DMi8 base and imaged using 20x or 40x objective lenses.

### Mycobacterium marinum

Mm infection experiments were performed using M. marinum M (ATCC #BAA-535), containing a psMT3-mCherry or psMT3 mCrimson vector (39). Injection inoculum was prepared from an overnight liquid culture in the log-phase of growth resuspended in 2% polyvinylpyrrolidone40 (PVP40) solution (CalBiochem) as previously described (40). 100 or 150 colony forming units (CFU) were injected into the caudal vein at 28-30hpf as previously described (41).

### Confocal microscopy of transgenic larvae

1dpi and 4dpi transgenic zebrafish larvae infected with fluorescent Mm strains were mounted in 0.8-1% low melting point agarose (Sigma-Aldrich) and imaged on a Leica TCS-SPE confocal on an inverted Leica DMi8 base and imaged using 20x or 40x objective lenses.

For quantification purposes acquisition settings and area of imaging were kept the same across groups. Corrected total cell fluorescence was calculated for each cell using Image J as previously described (42, 43).

### RNA injections

Embryos were injected with dominant *hif-1αb* (ZFIN: *hif1ab*) variant RNA at the one cell stage as previously described (44). *hif-1α* variants used were dominant active (DA) and dominant negative (DN) *hif-1α*. Phenol red (PR) (Sigma Aldrich) was used as a vehicle control.

### Pathway inhibitors

Unless otherwise stated, embryos were treated from 4 hours pre-Mm infection to 24 hours post infection (hpi) by addition to the embryo water and DMSO was used as a negative solvent control. The pan hydroxylase inhibitor, DMOG (dimethyloxaloylglycine, Enzo Life Sciences), was used at a 100μM concentration by incubation in E3 embryo media as previously described (44). The selective PHD inhibitors JNJ-402041935 (Cayman Chemicals) and FG-4592 (Selleckchem) were used at 100μM and 5μM respectively (45, 46). The selective inhibitor of COX-1, SC-560 (Cayman Chemicals), was used at 30μM and the selective inhibitor of COX-2, NS-398 (Cayman Chemicals), was used at 15μM by incubation in E3 embryo media as previously described (47). The 15-LOX inhibitor PD146176 (Tocris) was microinjected (1nl of 40nM) at 1 hour post infection (hpi). The leukotriene B4 receptor 1 (BLTR1) inhibitor, U-75302 (Abcam), was used at 30mM by incubation in E3 embryo media as previously described (48) together with the BLTR2 inhibitor, LY255283 (Abcam), which was used at 1mM. Exogenous prostaglandin E2 (Cayman Chemicals) and 15-keto prostaglandin E2 (Cayman Chemicals) were added by incubation in E3 at a concentration of 1μM (49).

### Hypoxia

Embryos were incubated in 5% oxygen (with 5% carbon dioxide) in a hypoxia hood (Baker-Ruskinn Sci-tive UM-027) from 32hpi for 6 hours and were imaged at 2dpf. Embryos from the same clutch kept in incubated normoxic room air were used as controls (33).

### Bacterial pixel count

Mm mCherry infected zebrafish larvae were imaged at 4dpi on an inverted Leica DMi8 with a 2.5x objective lens. Brightfield and fluorescent images were captured using a Hammamatsu OrcaV4 camera. Bacterial burden was assessed using dedicated pixel counting software as previously described (50).

### Human cells

Peripheral blood mononuclear cells (PBMC) were isolated by Ficoll-Paque PLUS (GE Healthcare) density centrifugation of whole blood from healthy donors (National Research Ethics Service reference 07/Q2305/7). PBMC were seeded at 2×10^6^/ml in 24-well tissue culture plates in RMPI 1640 media (Gibco) containing 2mmol/L L-glutamine (Lonza) and 10% newborn calf serum (Gibco). Cells were cultured in 5% CO_2_ at 37°C and non-adherent cells were removed after 24 hours. Remaining adherent cells were cultured for 14 days, with twice weekly media exchange, in RPMI 1640 supplemented with 2mmol/L L-glutamine and 10% heat-inactivated fetal bovine serum (FBS) (PAN Biotech) (51).

For experiments RPMI 1640 + 25mM HEPES (Gibco) containing 2mmol/L L-glutamine and 10% heat-inactivated FBS and drugs were pre-equilibrated in normoxia or hypoxia (0.8% O_2_, 5% CO_2_, 70% humidity at 37°C) for 24 hours prior to the experiment (52).

Cells were introduced to the hypoxic workstation (Baker-Ruskinn Sci-Tive UM-027) where culture media was removed and replaced with hypoxia-equilibrated media as detailed above. Cells were treated in duplicate or triplicate wells with 100µg/ml LPS (InvivoGen), DMSO (Sigma-Aldrich), or 15mM COX-2 inhibitor (NS398) (Cayman Chemical) and incubated for 18 hours. For normoxic controls, cells were treated identically within a class-2 tissue culture hood and transferred to a normoxic tissue-culture incubator.

Cell supernatants were collected following 18 hours incubation in normoxia or hypoxia and were frozen for subsequent analysis. Culture supernatants were assayed in duplicate or triplicate using a human TNF ELISA MAX (Biolegend). In samples where undiluted supernatants fell outside the standard curve, samples were re-run following 1:10 dilution in assay buffer.

### Statistical analysis

All data were analysed (Prism 7.0, GraphPad Software) using unpaired, two-tailed t-tests for comparisons between two groups and one-way ANOVA (with Bonferonni post-test adjustment) for other data. P values shown are: \**P* < .05, \*\**P* < .01, and \*\*\**P* < .001.

## Results

### *tnfa:GFP* expression is upregulated by injury, infection and Hif-1α stabilisation

TNF is a central regulator of the M1 inflammatory response to inflammation and pathogen challenge. Here, we use two transgenic zebrafish lines that express GFP via a *tnfa* promoter region; *TgBAC(tnfa:GFP)pd1028* and *Tg(tnfa:eGFP-F)ump5Tg* (31, 32). The *TgBAC(tnfa:GFP)pd1028* was made using BAC (bacterial artificial chromosome) transgenesis and contains 50kb of promoter region (32). The *Tg(tnfa:eGFP-F)ump5Tg* line has a smaller promoter region (3.8kb) than the BAC line (31) and has previously been demonstrated to be upregulated in macrophages after tailfin injury in a zebrafish model (31). Here, we demonstrate that this injury induced upregulation of macrophage *tnfa:GFP* is also found in the BAC transgenic line (*TgBAC(tnfa:GFP)pd1028* crossed to *Tg(mpeg1:mCherryCAAX)sh378)* shown 16 hours post wound (hpw) (Figure 1A). TNF has been implicated in all stages of TB infection, however the cell types involved *in vivo* have been difficult to observe (53). We used the *tnfa* BAC promoter GFP line to establish the transcriptional regulation of *tnfa* during pre- and early- larval Mm granuloma formation stages. *tnfa:GFP* expression was induced in larvae following Mm infection before granuloma onset (at 1 day post infection, dpi), and after granuloma formation (at 4dpi) (Figure 1B). *tnfa:GFP* expression was predominantly found in macrophages with internalised Mm, shown by co-localisation of fluorescence with *mpeg-1:mCherry* expressing macrophages and mCrimson expressing Mm (Figure 1C). Macrophage expression was confirmed using the other *tnfa* promoter driven line, (*Tg(tnfa:eGFP-F)ump5Tg*, crossed to *Tg(mpeg1:mCherry-F)ump2Tg*) (Figure S1). Our data demonstrate that injury and Mm induced *tnfa* expression occurs in macrophages as part of an early M1 response.

**Figure 1.**
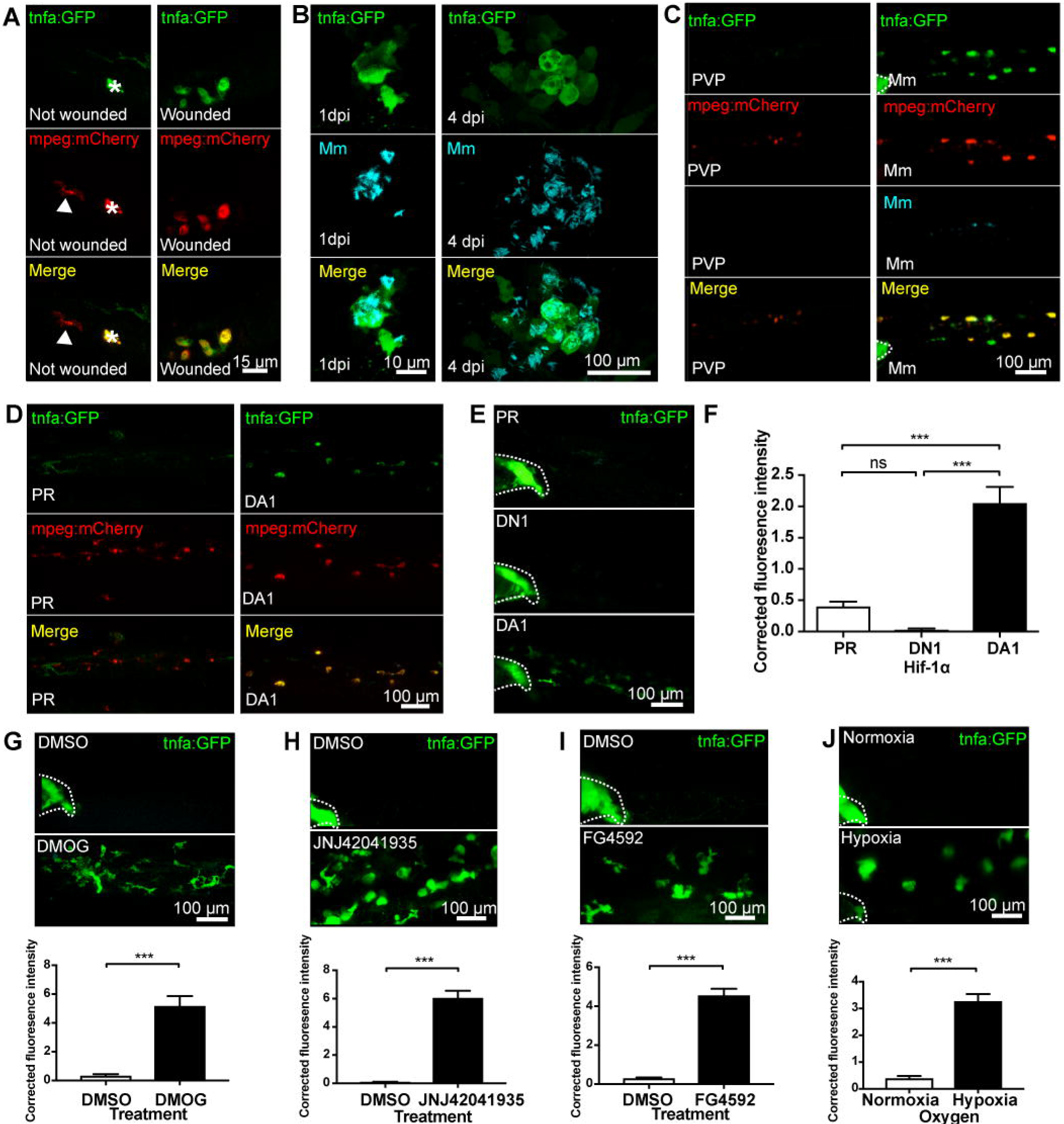
Macrophage *tnfa:GFP* is upregulated by injury, Mm infection and Hif-1 α stabilisation. (A) Fluorescent confocal micrographs of 3 days post fertilisation larvae with or without tailfin wound (induced 16 hours previously) with example macrophages at the tailfin site. *tnfa* expression was detected by GFP levels, in green, using the *TgBAC(tnfa:GFP)pd1028* transgenic line. Macrophages are shown in red using a *Tg(mpeg1:mCherryCAAX)sh378* line. The asterisk in the not wounded control indicates a neuromast that is fluorescent in all channels as a marker of position. (B) Fluorescent confocal micrographs of 1dpi Mm infected larvae, prior to granuloma formation (left panels), and 4 dpi, early granuloma (right panels) stages of infection. *tnfa* expression was detected by GFP levels, in green, using the *TgBAC(tnfa:GFP)pd1028* transgenic line. Mm mCrimson is shown in the far-red (cyan) channel. Increased levels of *tnfa:GFP* expression were detectable in cells associated with infection. (C) Fluorescent confocal micrographs of 1 day post infection (dpi) caudal vein region of infection. *tnfa* expression was detected by GFP levels, in green, using the *TgBAC(tnfa:GFP)pd1028* transgenic line. Macrophages are shown in red using a *Tg(mpeg1:mCherryCAAX)sh378* line. Mm Crimson is shown in the far-red channel (cyan, right panels) with a PVP control (left panels). Without infection there is little detectable *tnfa:GFP* expression, while infected and uninfected macrophages have higher levels of *tnfa:GFP* in the Mm group. Dotted lines indicate the yolk extension of the larvae where there is non-specific fluorescence. (D) Fluorescent confocal micrographs of 2 days post fertilisation larvae in the caudal vein region. *tnfa* expression was detected by GFP levels, in green, using the *TgBAC(tnfa:GFP)pd1028* transgenic line. Macrophages are shown in red using a *Tg(mpeg1:mCherryCAAX)sh378* line. Larvae were injected at the 1 cell stage with dominant active (DA) Hif-1α or phenol red (PR) control. (E) Fluorescent confocal micrographs of 2dpf zebrafish imaged around the caudal vein region. *tnfa* expression was detected by GFP levels, in green, using the *TgBAC(tnfa:GFP)pd1028* transgenic line. Larvae were injected at the 1 cell stage with dominant negative (DN) or dominant active (DA) Hif-1α or phenol red (PR) control. Dotted lines indicate the yolk extension of the larvae where there is non-specific fluorescence. (F) Graph shows corrected fluorescence intensity levels of *tnfa:GFP*. Dominant active Hif-1α (DA1, filled bar) had significantly increased *tnfa:GFP* levels compared to phenol red (PR) and dominant negative Hif-1α (DN1) injected controls (white bars). Data shown are mean ± SEM, n=66 cells from 12 embryos accumulated from 3 independent experiments and corresponds to images in (E). (G) Fluorescent confocal micrographs of 2dpf *TgBAC(tnfa:GFP)pd1028* transgenic larvae treated with DMOG or DMSO control. Dotted lines indicate the yolk extension of the larvae where there is non-specific fluorescence. Graph shows corrected fluorescence intensity levels of *tnfa:GFP* confocal z-stacks in larvae. DMOG treated larvae (filled bars) had significantly increased *tnfa:GFP* levels compared to DMSO controls (white bars). Data shown are mean ± SEM, n=24 cells from 6 embryos representative of 2 independent experiments. (G) Fluorescent confocal micrographs of 2dpf *TgBAC(tnfa:GFP)pd1028* transgenic larvae treated with the PHD inhibitor JNJ-42041935 or a DMSO solvent control. Dotted lines indicate the yolk extension of the larvae where there is non-specific fluorescence. Graph shows corrected fluorescence intensity levels of *tnf-a:GFP* confocal z-stacks at 2dpf treated with JNJ-42041935 (filled bars) or DMSO controls (white bars). Data shown are mean ± SEM, n=48 cells from 8 embryos accumulated from 2 independent experiments. (H) Fluorescent confocal micrographs of 2dpf *TgBAC(tnfa:GFP)pd1028* transgenic larvae treated with the PHD inhibitor FG4592 or a DMSO solvent control. Dotted lines indicate the yolk extension of the larvae where there is non-specific fluorescence. Graph shows corrected fluorescence intensity levels of *tnfa:GFP* confocal z-stacks in larvae at 2dpf treated with FG4592 (filled bars) or DMSO controls (white bars). Data shown are mean ± SEM, n=54 cells from 9 embryos accumulated from 3 independent experiments. (I) Fluorescent confocal micrographs of 2dpf *TgBAC(tnfa:GFP)pd1028* transgenic larvae incubated in room normoxia or 5% oxygen (hypoxia) for 6 hours between 32-38hpf. Dotted lines indicate the yolk extension of the larvae where there is non-specific fluorescence. Graph shows corrected fluorescence intensity levels of *tnfa:GFP* confocal z-stacks in larvae at 2dpf incubated in 5% oxygen (filled bars) or normoxia controls (white bars). Data shown are mean ± SEM, n=72 cells from 12 embryos accumulated from 3 independent experiments.

Hif-1α can be stabilised in zebrafish both genetically and pharmacologically using dominant active constructs and hydroxylase inhibitors respectively (29, 44). We tested whether Hif-1α promotes M1 polarisation using the *tnfa:GFP* BAC transgenic line as a readout of M1 macrophages (31, 32). Hif-1α was stabilised genetically in larvae using DA Hif-1α RNA (44), which resulted in upregulation of *tnfa:GFP* expression (Figure 1D-F) indicating a shift of macrophage phenotype towards an activated M1 response. Dominant negative (DN) Hif-1α caused no upregulation of *tnfa:GFP* expression (Figure 1E-F). To demonstrate that it was also possible to boost the Hif-1α M1 pro-inflammatory response pharmacologically, the hypoxia mimetic DMOG (dimethyloxaloylglycine, a pan-hydroxylase inhibitor that stabilises Hif-1α via inactivation of regulatory prolyl hydroxylase enzymes) was used to stabilise endogenous levels of Hif-1α (57). DMOG treatment upregulated expression of *tnfa*:*GFP* compared to DMSO solvent controls, phenocopying the DA-Hif-1α response (Figure 1G). Recently described hypoxia mimetics with reportedly greater prolyl hydroxylase selectively, JNJ-42041935 and FG4592, had similar effects to DMOG (Figure 1H and I) (33, 45, 46). During both homeostasis and disease physiology, Hif-1α stability is regulated by microenvironmental hypoxia. To simulate this we subjected the *tnfa:GFP* line to 5% oxygen for 6 hours at 32hpf and looked for GFP expression at 48hpf. This level of hypoxia was sufficient to switch on GFP expression in the *Tg(phd3:GFP)i144* hypoxia reporter zebrafish (Figure S2) (33). Six hours of 5% oxygen increased *tnfa:GFP* expression at 48hpf, compared to normoxic controls, to a similar level as that observed with genetic or pharmacological Hif-1α stabilisation (Figure 1J). Together, these data indicate that *tnfa* expression is part of a pro-inflammatory M1 macrophage response to hypoxia and stabilised Hif-1α, a response that is targetable by pharmacological agents and has the potential to aid the host response to bacterial challenge.

### Hif-1α dependent *tnfa:GFP* transcription requires cyclooxygenase

Eicosanoids are lipid signalling molecules produced by macrophages during early microbial infection as part of an M1 response (22, 23). We tested whether cyclooxygenase inhibition affected Hif-1α mediated *tnfa:GFP* production. At basal levels, *tnfa:GFP* was not altered by the Cox-1 inhibitor SC560 or the Cox-2 inhibitor NS398 compared to DMSO treated negative controls (Figure 2A-B). Strikingly, DA Hif-1α-induced *tnfa:GFP* expression was significantly abrogated to basal levels by both SC560 and NS398 (Figure 2A-B). We have previously shown that DA Hif-1α induces the expression of another important pro-inflammatory cytokine, *il-1β*, shown by an *il-1β:GFP* transgenic line (Figure S3A) (33). Cox-1 inhibition by SC560 did not abrogate Hif-1α induced *il-1β:GFP*, while Cox-2 inhibition with NS398 caused a small decrease that did not reach basal levels (Figure S3B-C). These data indicate that Hif-1α-induced *tnfa* expression is caused by a product of the cyclooxygenase arm of the arachidonic acid pathway via a novel macrophage HIF/COX/TNF axis (Figure 2C), while Hif-1α induced *il-1β* does not fully act via this pathway suggestive of complex regulation of M1 pro-inflammatory cytokines by Hif-1α stabilisation.

**Figure 2.**
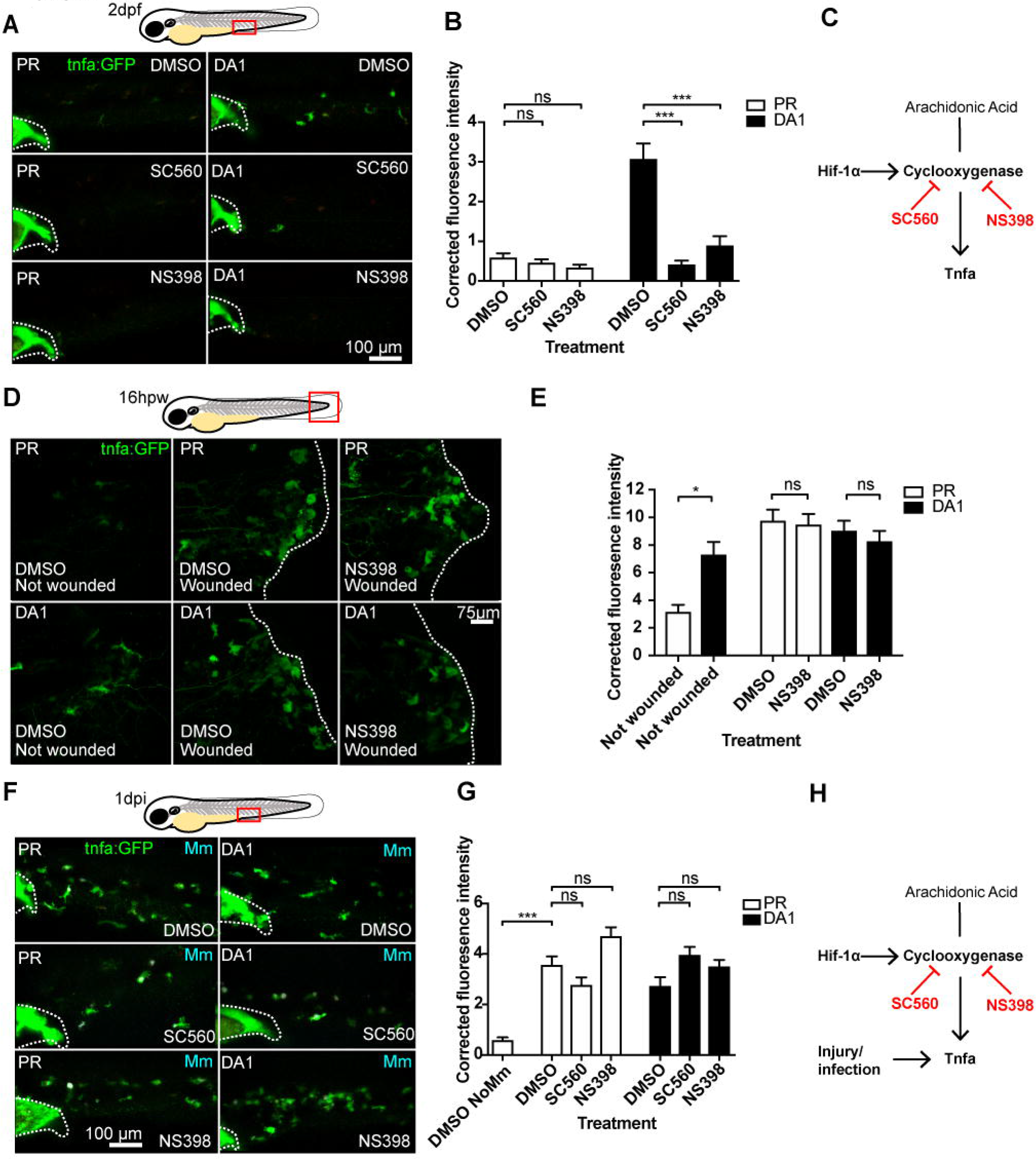
Hif-1α-activated *tnfa:GFP* is cyclooxygenase dependent while injury and infection induced *tnfa:GFP* is cyclooxygenase independent. (A) Fluorescent confocal micrographs of 2dpf caudal vein region of larvae. *tnfa* expression was detected by GFP levels, in green, using the *TgBAC(tnfa:GFP)pd1028* transgenic line. Phenol red (PR) and dominant active Hif-1α (DA1) injected larvae were treated with DMSO, SC560 (Cox-1 inhibitor) and NS398 (Cox-2 inhibitor). Dotted lines indicate the yolk extension of the larvae where there is non-specific fluorescence. (B) Corrected fluorescence intensity levels of *tnfa:GFP* confocal z-stacks in PR (white bars) and DA1 (filled bars) injected larvae at 2dpf treated with Cox inhibitors. Data shown are mean ± SEM, n=36 cells from 6 embryos representative of 3 independent experiments. (C) Schematic of the arachidonic pathway indicating where stabilising Hif-1α upregulates *tnfa* via cyclooxygenase, an effect that is blocked using the Cox1/2 inhibitors SC560/NS398. (D) Fluorescent confocal micrographs of 3 days post fertilisation larvae with or without tailfin wound (induced 16 hours previously) in phenol red (PR) or dominant active Hif-1α embryos. *tnfa* expression was detected by GFP levels, in green, using the *TgBAC(tnfα:GFP)pd1028* transgenic line. Embryos were treated with DMSO or NS398 (Cox-2 inhibitor). (E) Corrected fluorescence intensity levels of *tnfa:GFP* confocal z-stacks in PR (white bars) and DA1 (filled bars) injected larvae at 2dpf treated with Cox inhibitors. Data shown are mean ± SEM, n=24 (in not wounded) or n=36 (in wounded) cells from 6 embryos accumulated from 2 independent experiments. Not-wounded fish had fewer macrophages at the tailfin site, hence the lower cell number in these groups. (F) Fluorescent confocal micrographs of 1dpi caudal vein region of Mm Crimson infected larvae. *tnfa* expression was detected by GFP levels, in green, using the *TgBAC(tnfa:GFP)pd1028* transgenic line. Phenol red (PR) and dominant active Hif-1α (DA1) injected larvae were treated with DMSO, SC560 (Cox-1 inhibitor) and NS398 (Cox-2 inhibitor) and infected with Mm Crimson. Dotted lines indicate the yolk extension of the larvae where there is non-specific fluorescence. (G) Corrected fluorescence intensity levels of *tnfa:GFP* confocal z-stacks in PR (white bars) and DA1 (filled bars) Mm Crimson infected larvae at 1dpi treated with Cox inhibitors. NB, DMSO NoMm is a no Mm control taken from figure 2B for comparison with Mm infected data (data is from the same experiment). Data shown are mean ± SEM, n=36 cells from 6 embryos representative of 3 independent experiments. (H) Schematic showing that injury and infection upregulate *tnfa* independently of cyclooxygenase, an effect that is not blocked by the Cox1/2 inhibitors SC560/NS398.

We next tested whether injury- and infection-induced *tnfa* are cyclooxygenase dependent processes. Macrophage *tnfa:GFP* expression induced after injury was not abrogated by cyclooxygenase inhibition using NS398 (Figure 2D-E). Similarly, cyclooxygenase inhibition using either SC560 or NS398 did not alter the expression of Mm-induced *tnfa:GFP* (Figure 2F-G). Our data indicate that macrophage *tnfa* expression can be driven by the presence of DAMPS (injury) or PAMPS (Mm) and that these are cyclooxygenase independent mechanisms (Figure 2H).

We have previously shown that Hif-1α stabilisation is host-protective in the zebrafish Mm model, an effect that is dependent on macrophage expression of proinflammatory *il-1β* (33). We therefore hypothesised that priming macrophages with increased *tnfa* via Hif-1α stabilisation may be protective against subsequent Mm infection. Neither genetic nor pharmacological Hif-1α stabilisation were additive to the *tnfa:GFP* expression caused by either injury or Mm infection (Figure 2C-F and S4A-D). To test whether DA Hif-1α priming of macrophage *tnfa* had an effect on the outcome of Mm infection, bacterial burden was quantified in DA Hif-1α-injected embryos with cyclooxygenase inhibition (having demonstrated that Hif-1α-induced *tnfa* requires cyclooxygenase activity in Figure 2A-C). Neither SC560 nor NS398 mediated inhibition of Hif-1α-induced *tnfa* expression diminished the host-protective effect of DA Hif-1α (Figure S5A-C) or DMOG treated larvae (Figure S5D-F). Taken together these data indicate that injury- and Mm- induced *tnfa* is not further increased by Hif-1α stabilisation and that cyclooxygenase mediated early priming of *tnfa* in macrophages is not required for the Hif-1α mediated reduction of bacterial burden.

### Blocking cyclooxygenase independent arachidonic acid pathways does not abrogate Hif-1α upregulation of *tnfa:GFP*

To investigate whether the effect of the cyclooxygenase inhibitors on *tnfa* was specific to the prostaglandin path of the arachidonic acid pathway, we targeted the lipoxin and leukotriene producing arms using the 15-Lipoxygenase inhibitor PD146176 and leukotriene B4 receptor antagonists. The 15-Lipoxygenase inhibitor PD146176 did not block the *tnfa:GFP* induced by DA Hif-1α (Figure 3A-B). Furthermore, PD147176 increased *tnfa:GFP* expression on its own in the absence of infection, although not to the same extent as DA Hif-α (Figure 3A-B). PD146176 also did not affect *tnfa:GFP* expression after Mm infection (Figure 3A-B), nor did it block the protective effect of DA Hif-1α. Treatment with PD146176 was sufficient to decrease Mm burden, but not to the same extent as DA Hif-1α (Figure 3C). Leukotriene B4 inhibition, using the BLTR1/2 antagonists U75302 and LY255283, did not increase *tnfa:GFP* levels and also did not block DA Hif-1α induced *tnfa:GFP* (Figure 3D-E). These data indicate that blocking components of the lipoxygenase dependent arms of the arachidonic acid pathway do not block the Hif-1α effect on *tnfa* expression and do not replicate the effects observed by blocking the cyclooxygenase dependent arm (Figure 3F).

**Figure 3.**
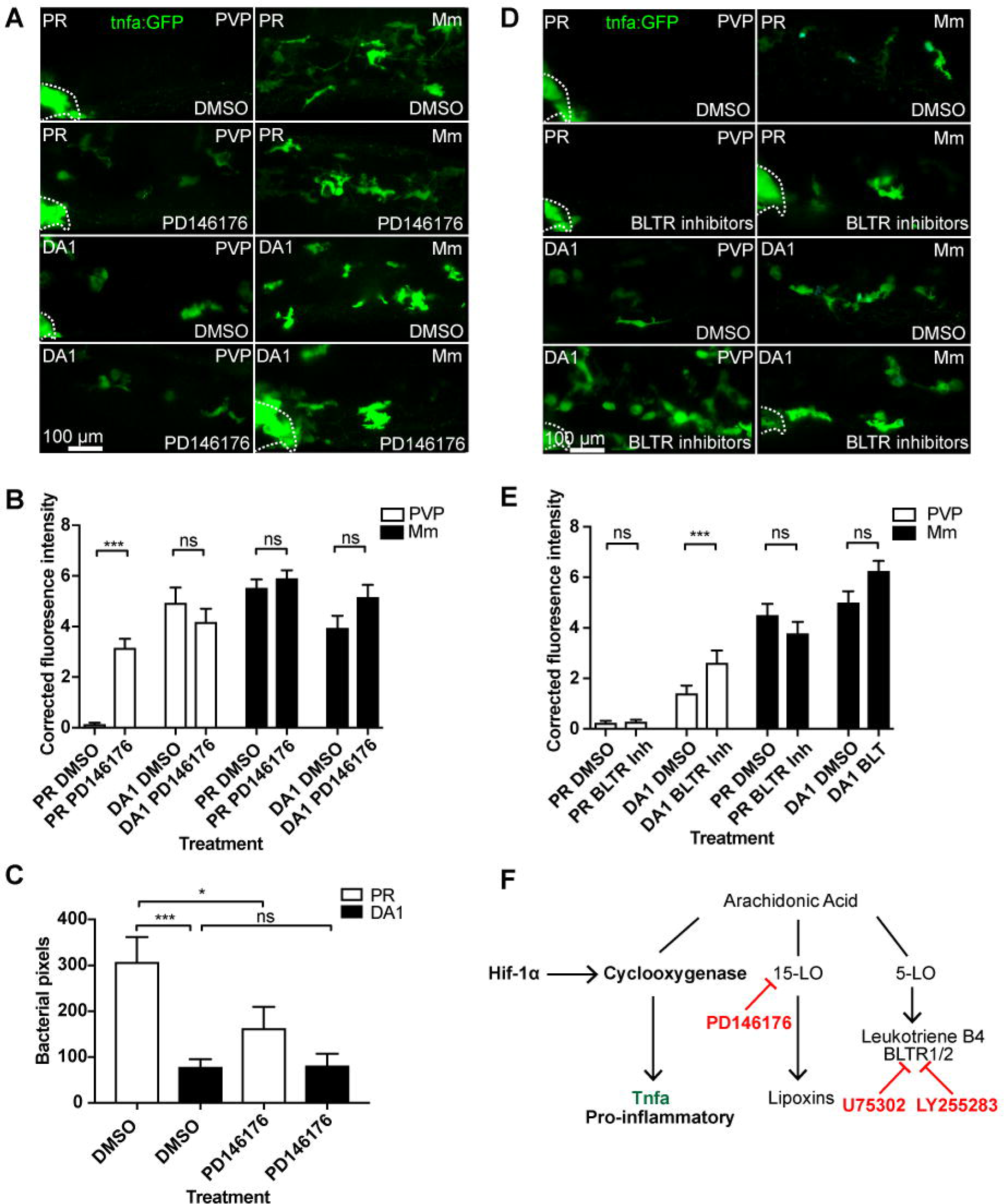
Blocking 15-lipoxygenase or leukotriene B4 receptors does not abrogate DA-Hif-1α-upregulation of *tnfa:GFP*. (A) Fluorescent confocal micrographs of 1dpi caudal vein region of Mm and PVP infected larvae. *tnfa* expression was detected by GFP levels, in green, using the *TgBAC(tnfa:GFP)pd1028* transgenic line. Phenol red (PR) and dominant active Hif-1α (DA1) injected larvae were treated with DMSO or PD146176 (15-Lipoxygenase inhibitor). Non-infected larvae are in the left panels (PVP) and Mm Crimson infected larvae are in the right panels (Mm). Dotted lines indicate the yolk extension of the larvae where there is non-specific fluorescence. (B) Corrected fluorescence intensity levels of *tnfa:GFP* confocal z-stacks in uninfected larvae (PVP, empty bars) and infected larvae (Mm, filled bars) at 1dpi of data shown in (A) after treatment with DMSO or PD146176 (15-Lipoxygenase inhibitor). Data shown are mean ± SEM, n=42 cells from 7 embryos accumulated from 3 independent experiments. (C) Bacterial burden at 4dpi after injection of DA Hif-1α (DA1) and 24 hours of the 15-lipoxygenase inhibitor PD146176, using DMSO as a negative solvent control. Data shown are mean ± SEM, n=48-50 as accumulated from 3 independent experiments. (D) Fluorescent confocal micrographs of 1dpi caudal vein region of Mm and PVP infected larvae shown in A. *tnfa* expression was detected by GFP levels, in green, using the *TgBAC(tnfa:GFP)pd1028* transgenic line. Phenol red (PR) and dominant active Hif-1α (DA1) injected larvae were treated with DMSO or U75302/LY255283 (BLTR1/2 inhibitors). Non-infected larvae are in the left panels (PVP) and Mm Crimson infected larvae are in the right panels (Mm). Dotted lines indicate the yolk extension of the larvae where there is non-specific fluorescence. (E) Corrected fluorescence intensity levels of *tnfa:GFP* confocal z-stacks in uninfected larvae (PVP, empty bars) and infected larvae (Mm, filled bars) at 1dpi of data shown in (D) after treatment with DMSO or U75302/LY255283 (BLTR1/2 inhibitors). Data shown are mean ± SEM, n=30 cells from 5 embryos accumulated from 2 independent experiments. (F) Schematic of the arachidonic pathway indicating where stabilising Hif-1α upregulates tnfa via cyclooxygenase, an effect that is not blocked using the 15-lipoxygenase inhibitor PD146176 or BLTR1/2 inhibitors U75302/LY255283.

### Hif-1α-induced *tnfa:GFP* requires active prostaglandin E2

A key family of immune regulators downstream of arachidonic acid and cyclooxygenases are prostaglandins. The best characterised of these as a regulator of macrophage function is prostaglandin E2 (PGE2). We tested whether PGE2 was a mediator in the HIF/COX/TNF pathway by addition of exogenous PGE2 to DA Hif-1α larvae. Exogenous PGE2 had no effect on *tnfa:GFP* expression in the absence or presence of Mm infection (Figure 4A-D). However, PGE2 was able to rescue the decrease in *tnfa:GFP* expression after Cox-1 (Figure 4A and C) or Cox-2 (Figure 4B and D) inhibition in the DA Hif-1α larvae. Furthermore, addition of PGE2 alone led to a significant increase in the DA Hif-1α-induced *tnfa:GFP* expression (Figure 4A-D). These rescuing effects were not observed by addition of exogenous 15-keto prostaglandin E2, an immunologically inactive degradation product of PGE2 (Figure 4E-F) (49, 58, 59). These data indicate that Hif-1α-induced *tnfa:GFP* requires active prostaglandin E2 (Figure 4G).

**Figure 4.**
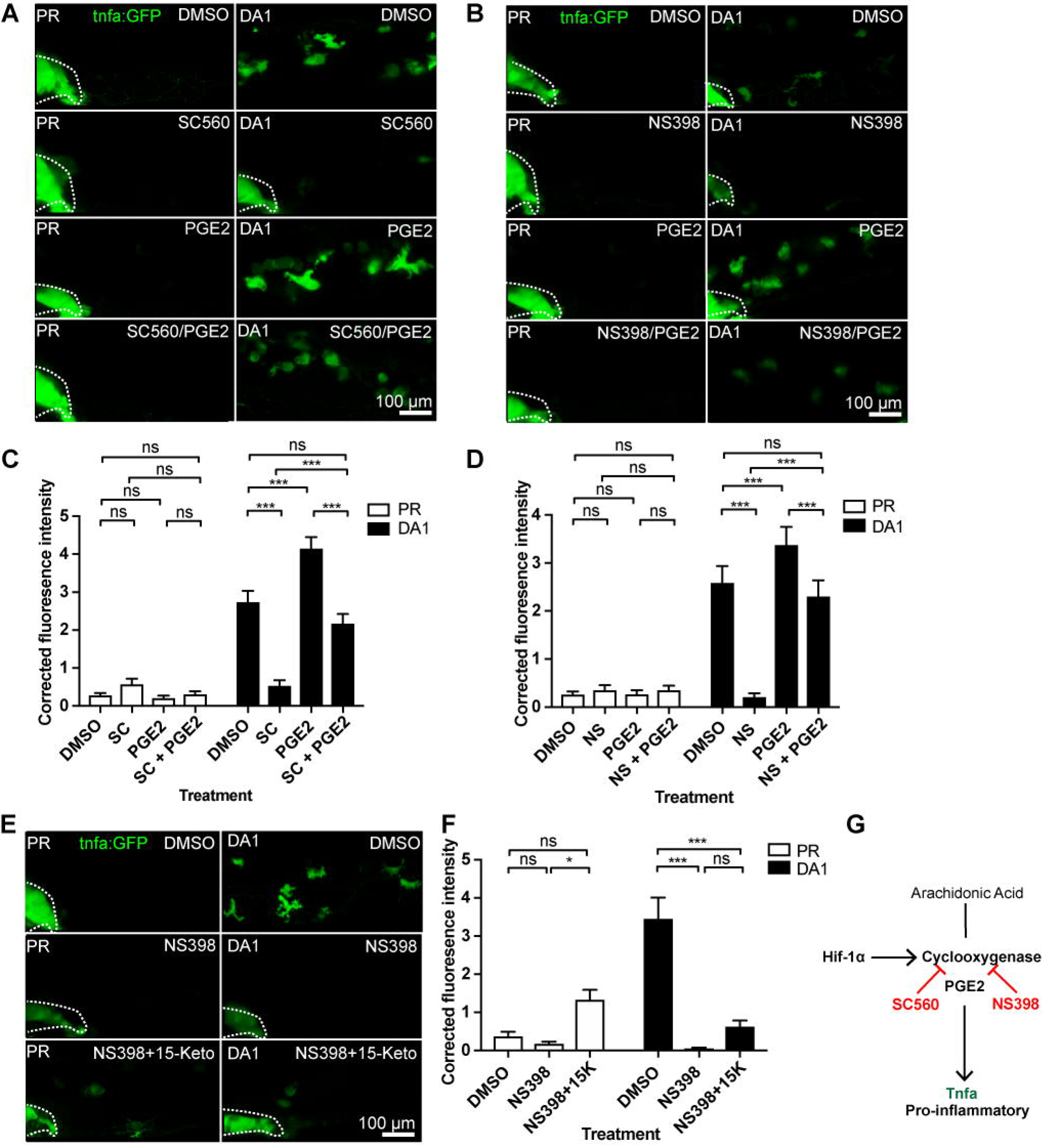
Hif-1α-induced *tnfa:GFP* requires active prostaglandin E2. (A) Fluorescent confocal micrographs of 1dpi caudal vein region of Mm. *tnfa* expression was detected by GFP levels, in green, using the *TgBAC(tnfa:GFP)pd1028* transgenic line. Phenol red (PR) and dominant active Hif-1α (DA1) injected larvae were treated with DMSO or SC560 (Cox-1 inhibitor) in the presence or absence of endogenous prostaglandin E2 (PGE2). All larvae are PVP injected. Dotted lines indicate the yolk extension of the larvae where there is non-specific fluorescence. (B) Fluorescent confocal micrographs of 1dpi caudal vein region. *tnfa* expression was detected by GFP levels, in green, using the *TgBAC(tnfa:GFP)pd1028* transgenic line. Phenol red (PR) and dominant active Hif-1α (DA1) injected larvae were treated with DMSO or NS398 (Cox-2 inhibitor) in the presence or absence of endogenous prostaglandin E2 (PGE2). All larvae are PVP injected. Dotted lines indicate the yolk extension of the larvae where there is non-specific fluorescence. (C) Corrected fluorescence intensity levels of *tnfa:GFP* confocal z-stacks in PVP injected phenol red (PR, empty bars) and dominant active Hif-1α (DA1, filled bars) larvae at 1dpi of data shown in (B) after treatment with DMSO/NS398/PGE2. Data shown are mean ± SEM, n=54 cells from 9 embryos accumulated from 3 independent experiments. (D) Corrected fluorescence intensity levels of *tnfa:GFP* confocal z-stacks in PVP injected phenol red (PR, empty bars) and dominant active Hif-1α (DA1, filled bars) larvae at 1dpi of data shown in (B) after treatment with DMSO/NS398/PGE2. Data shown are mean ± SEM, n=54 cells from 9 embryos accumulated from 3 independent experiments. (E) Fluorescent confocal micrographs of 1dpi caudal vein region. *tnf-a* expression was detected by GFP levels, in green, using the *TgBAC(tnfa:GFP)pd1028* transgenic line. Phenol red (PR) and dominant active Hif-1α (DA1) injected larvae were treated with DMSO or NS398 (Cox-2 inhibitor) in the presence or absence of endogenous 15-keto prostaglandin E2 (15K). All larvae are PVP injected. Dotted lines indicate the yolk extension of the larvae where there is non-specific fluorescence. (F) Corrected fluorescence intensity levels of *tnfa:GFP* confocal z-stacks in PVP injected phenol red (PR, empty bars) and dominant active Hif-1α (DA1, filled bars) larvae at 1dpi of data shown in (E) after treatment with DMSO/NS398/15-keto PGE2. Data shown are mean ± SEM, n=24 cells from 4 embryos accumulated from 2 independent experiments. (G) Schematic of the arachidonic pathway indicating where blocking cyclooxygenase prevents Hif-1α upregulation of *tnfa*, an effect rescued by endogenous PGE2.

### The HIF-COX-TNF axis is conserved in human macrophages

To translate our findings from zebrafish to humans we tested whether HIF-1α stabilisation in human macrophages induces TNF expression. We found that human monocyte derived macrophages (hMDMs) produced higher levels of TNF protein in hypoxia (0.8% oxygen) than those in normoxia, measured by an anti-human TNF ELISA (Figure 5A). Furthermore, treatment with the COX-2 inhibitor, NS398, reduced this hypoxia-induced TNF back to the equivalent levels found in normoxia (Figure 5A). This was replicated when HIF-1α was stabilised in hHDMs using the hypoxia mimetic FG4592 (Figure 5B). These data indicate that the HIF/COX/TNF axis is a conserved mechanism in human macrophages and could be important in human disease.

**Figure 5.**
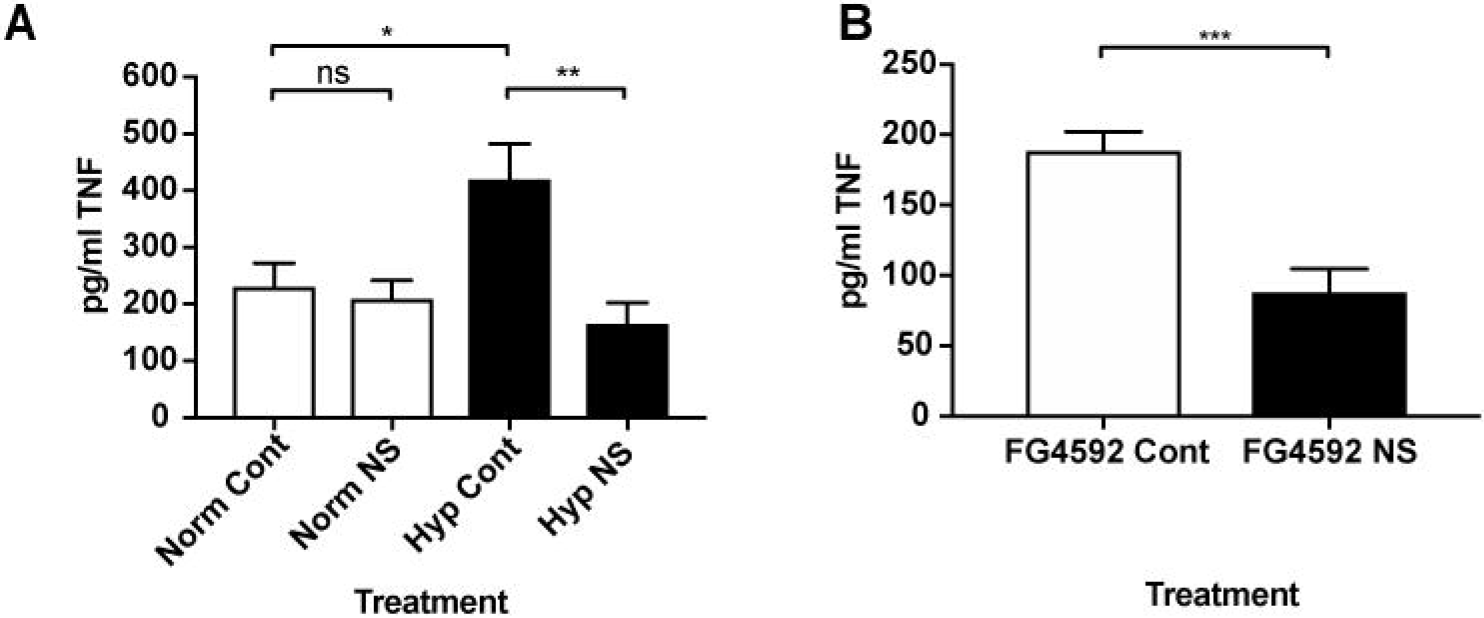
Hypoxia induces TNF in human MDMs in a COX-2 dependent manner. (A) TNF ELISA of human monocyte derived macrophages incubated in normoxia or 0.8% hypoxia with or without treatment with NS398. Data shown are mean ± SEM, n=5-8 biological repeats from 3-4 donors. (B) TNF ELISA of human monocyte derived macrophages incubated in normoxia treated with FG4592 with or without treatment with NS398. Data shown are mean ± SEM, n=6 biological repeats from 3 donors.

## Discussion

Control of macrophage function during homeostasis and infection are critical for healthy tissues and must integrate changes in the local microenvironment with cytokine signalling. Understanding of signalling pathways that link the microenvironment with macrophage phenotypic outcomes will identify potential novel therapeutic avenues for control of macrophages during disease. Currently, the mechanisms and molecular cues leading to different macrophage phenotypes are not well-defined *in vivo*.

Here we show that a disease relevant microenvironmental cue, hypoxia signalling via Hif-1α, upregulates macrophage *tnfa* expression in a cyclooxygenase dependent manner *in vivo*. TNF regulation by hypoxia has been shown in a range of mammalian cells, and its promoter region contains HIF responsive elements (HREs), resulting in some level of direct regulation by HIF-1α (60–63). We observed an M1 pro-inflammatory *tnfa* response with stabilised Hif-1α, demonstrating that hypoxia signalling alone can lead to a switch of macrophage polarisation to an M1 phenotype, consistent with our previous observations concerning *il-1β* activation (33). Not only could *tnfa* activation be achieved by genetic stabilisation of Hif-1α, but also using hypoxia mimetics and physiological hypoxia. These findings indicate that the Hif-1α pro-inflammatory switch is targetable by pharmaceuticals and could be druggable during disease. Hypoxic regulation of TNF via COX has previously been demonstrated in mammalian osteoblasts, however, little is known about this interaction (26). Our data demonstrate that Hif-1α upregulation of macrophage *tnfa* is dependent on cyclooxygenase and further shows that the mechanism is likely to be via the production of PGE2. The degradation metabolite 15-keto-PGE2 did not rescue the loss of *tnfa* expression after cyclooxygenase inhibition, consistent with previous reports that 15-keto-PGE2 is unable to bind the prostaglandin E2 EP receptors, demonstrating a requirement for active PGE2 (64).

Mycobacteria mediated macrophage polarisation is complex, with M1 and M2 phenotypes observed during pathogenesis, however, the mechanisms determining macrophage phenotype are not fully characterised (56). The Mtb/Mm granuloma has been widely studied in human, mice and zebrafish and is rich in pro-inflammatory cytokine production (32, 33, 65–69). Here, we observed that an M1 pro-inflammatory *tnfa* response was induced at pre-granuloma stages as well as during granuloma formation *in vivo*. A key advantage of using fluorescent transgenic zebrafish lines, such as *tnfa:GFP*, is that they enable identification of the cell type producing the transgene in an intact organism. This is especially important when studying early innate immune cell interactions with pathogens, as it eliminates the risk of activation during a sorting process. TNF is required for the control of initial TB infection in mice and also of latent TB in humans (68, 70–73). Patients on anti-TNF therapies for immune diseases such as Crohn’s disease and rheumatoid arthritis, which have proved effective treatments for these debilitating illnesses, have an associated increased risk of TB reactivation and infectious disease (72, 74). In the zebrafish Mm model it has been widely demonstrated that Tnfa is required for control of early infection, with perturbation of Tnfa signalling leading to high infectious burdens (75). When Hif-1α driven *tnfa* was downregulated by cyclooxygenase inhibition, there was no effect on the decrease in bacterial burden resulting from stabilisation of Hif-1α. Furthermore, none of the arachidonic acid pathway inhibitors used in our study had any effect on Tnfa production after Mm infection. These findings are consistent with Mm induced macrophage Tnfa production being independent of arachidonic acid, and is most likely TLR mediated as has been reported elsewhere (43, 56). Our data indicate that there is no additional protective effect of priming macrophages with increased levels of Tnfa before infection onset. Interestingly, this is not the case for another important M1 cytokine, Il-1β, where Hif-1α stabilisation fails to reduce bacterial burden if Il-1β is blocked early in infection, (33). Our findings indicate that different M1 cytokines induced by Hif-1α stabilisation can have differential effects on the outcome of infection, and highlight that, although TNF and IL-1β are often used as markers of M1 polarisation, their roles in the pro-inflammatory macrophage are distinct.

Blocking cyclooxygenase independent arms of the arachidonic acid pathway did not inhibit Hif-1α-induced macrophage *tnfa* transcription in the same way as blocking the cyclooxygenase/PGE2 arm. Inhibition of the 15-lipoxygenase arm was sufficient to induce macrophage *tnfa* transcription which was host protective and decreased Mm burden. These findings indicate a potential shift towards the COX/TNF dependent axis when 15-lipoxygenase is blocked and are consistent with previous observations suggesting that inhibition of 15-Lipoxygenase can have immune activating effects (48).

Our data reveal a novel mechanism of TNF activation in macrophages via a Hif-1α/cyclooxygenase/PGE2 axis. Importantly, this pathway is conserved in human macrophages, meaning that it could be targetable for pharmaceutical intervention to promote macrophage M1-type polarisation in human disease. Although the effect of the HIF/COX/TNF axis in early Mm infection was negligible, this pathway is likely to have important roles in numerous disease situations. Multiple disease pathologies have hypoxia as a features of the tissue microenvironment, concurrent with macrophage influx. Hif-1α mediated M1 switching and TNF activation may be pertinent in later TB infection situations where granulomas have stabilised Hif-1α due to necrotic and hypoxic centres (76, 77). It is not only bacterial infections such as TB that might be influenced by a HIF-1α mediated M1/TNF switch. Hypoxia is a key hallmark of cancers with high levels of HIF-1α widely found in those that produce large tumours where the centre is hypoxic (78). This is also true of cyclooxygenase/PGE2 production, where these important immunomodulatory signals are required for cellular adaption to the tumour microenvironment (79). Certain types of cancer induce inflammatory microenvironments, where cytokines and other immunomodulators can contribute to the progression of cancers (80). Macrophages play central roles in cancer-inflammation that could be exploited utilising pharmaceutical control of this novel HIF/COX/TNF mechanism. The zebrafish larvae is small enough to be fully oxygenated at the 2-5dpf stages of this study (81). Further investigation of the HIF/COX/TNF axis in models where hypoxia is a key hallmark of disease pathology is required to uncover the full therapeutic relevance of this potentially important novel macrophage pathway.

In conclusion, we have identified a novel mechanism for macrophage *tnfa* transcription via Hif-1α and cyclooxygenase that is conserved from zebrafish to humans, highlighting a druggable mechanism to induce an M1-macrophage response. Importantly, this axis links a microenvironmental cue, hypoxia, to a macrophage pro-inflammatory M1 cytokine, *tnfa* expression. We provide strong evidence to show that this response acts via the cyclooxygenase/PGE2 arm of the arachidonic pathway. Due to the central roles of these modulators in disease microenvironments we anticipate that this HIF/COX/TNF pathway may have important implications in conditions such as inflammation, infection and cancer.

## Acknowledgements

The authors would like to thank The Bateson Aquarium Team for fish care and the IICD Technical Team for practical assistance (University of Sheffield). We gratefully thank, Michel Bagnat (Duke University) for providing their *TgBAC(tnfa:GFP)pd1028* transgenic line, Georges Lutfalla (Montpellier University) for providing the *Tg(tnfa:eGFP-F)ump5Tg* line, Annemarie Meijer (University of Leiden) for M. marinum strains. We thank David Dockrell (University of Edinburgh), Paul Collini (University of Sheffield) and Jon Kilby (University of Sheffield) for help with hMDM experiments. Thanks to Alison Condliffe and Benjamin Durham (University of Sheffield) for use of, and invaluable help with, their hypoxia hood. We would like to thank Robert Evans (The Francis Crick, London) and Catherine Loynes (University of Sheffield) for helpful discussions and advice on prostaglandin modulation. Thanks to Simon Johnston, Hannah Isles and Stephen Renshaw (University of Sheffield) for constructive comments on the manuscript.

## Author Contributions

Conceived and designed the experiments: PME, AL. Performed the experiments: PME, AL. Analysed the data: PME, AL. Wrote the paper: PME.

## Figure Legends

**Figure S1. Mm infection induced *tnfa:GFP* in macrophages in a second transgenic line.**

Fluorescent confocal micrographs of 1dpi caudal vein region of infection. *tnfa* expression was detected by GFP levels, in green, using the *Tg(tnfa:eGFP-F)ump5Tg* transgenic line. Macrophages are shown in red using a *Tg(mpeg1:mCherry-F)ump2Tg* line. Mm Crimson is shown in the blue channel. Dotted lines indicate the yolk extension of the larvae where there is non-specific fluorescence.

**Figure S2. 5% oxygen induces expression of the *phd3:GFP* hypoxia transgene.**

Fluorescent micrographs of 48hpf *Tg(phd3:eGFP)i144* hypoxia reporter zebrafish after incubated in 6 hours of 5% oxygen from 32-38hpf, or normoxic controls.

**Figure S3. DA-Hif-1α induced *il-1β:GFP* is not altered by Cox-1 inhibition and decreased by Cox-2 inhibition.**

(A) Fluorescent confocal micrographs of 2dpf caudal vein region. *il-1β* expression was detected by GFP levels, in green, using the *TgBAC(il-1β:GFP)SH445* transgenic line. Macrophages are shown in red using a *Tg(mpeg1:mCherryCAAX)sh378* line. Without infection there is little detectable *il-1β:GFP* expression in phenol red (PR) controls, while DA-Hif-1α macrophages have higher levels of *il-1β:GFP* in the Mm group.

(B) Fluorescent confocal micrographs of 1dpi caudal vein region of PVP injected larvae. *il-1β* expression was detected by GFP levels. Phenol red (PR) and dominant active Hif-1α (DA1) injected larvae were treated with DMSO, SC560 (COX-1 inhibitor) and NS398 (COX-2 inhibitor).

(C) Corrected fluorescence intensity levels of *il-1β:GFP* confocal z-stacks in PVP injected larvae at 1dpi from (B). Data shown are mean ± SEM, n=36 cells from 6 embryos representative of 3 independent experiments.

**Figure S4. Mm infection induced *tnfa:GFP* is not altered by Hif-1α stabilisation**

(A) Fluorescent confocal micrographs of Mm Crimson infected 1dpi zebrafish imaged around the caudal vein region. *tnfa* expression was detected by GFP levels, in green, using the *TgBAC(tnfa:GFP)pd1028* transgenic line. Larvae were injected at the 1 cell stage with dominant negative (DN) or dominant active (DA) Hif-1α or phenol red (PR) control. Dotted lines indicate the yolk extension of the larvae where there is non-specific fluorescence.

(B) Graph shows corrected fluorescence intensity levels of *tnfa:GFP*. Dominant active Hif-1α (DA1, filled bar) showed no change in *tnfa:GFP* levels in the presence of Mm bacterial challenge compared to phenol red (PR) and dominant negative Hif-1α (DN1) injected controls (white bars). NB, PR NoMm is a no Mm control taken from figure 1A for comparison with Mm infected data, the data for which is from the same experiment as this one. Data shown are mean ± SEM, n=66 cells from 12 embryos accumulated from 3 independent experiments.

(C) Fluorescent confocal micrographs of 2dpf Mm infected *TgBAC(tnfa:GFP)pd1028* transgenic larvae treated with DMOG or DMSO control. Dotted lines indicate the yolk extension of the larvae where there is non-specific fluorescence.

(D) Graph shows corrected fluorescence intensity levels of *tnfa:GFP* confocal z-stacks in larvae. DMOG treated larvae (filled bars) had significantly increased *tnfa:GFP* levels compared to DMSO controls (white bars). Data shown are mean ± SEM, n=24 cells from 6 embryos representative of 2 independent experiments.

**Figure S5. Cyclooxygenase inhibition does not block the protective effect of Hif-1α stabilisation in Mm infection.**

(A) Bacterial burden of SC560 treated larvae at 4dpi after injection with phenol red (PR) or dominant active Hif-1α (DA1). Data shown are mean ± SEM, n=35 as accumulated from 3 independent experiments.

(B) Bacterial burden of NS398 treated larvae at 4dpi after injection with phenol red (PR) or dominant active Hif-1α (DA1). Data shown are mean ± SEM, n=35 as accumulated from 3 independent experiments.

(C) Bacterial burden of DMSO treated larvae at 4dpi after co-treatment with DMSO or DMOG. Data shown are mean ± SEM, n=33-36 as accumulated from 3 independent experiments.

(D) Bacterial burden of SC560 treated larvae at 4dpi after co-treatment with DMSO or DMOG. Data shown are mean ± SEM, n=33 as accumulated from 3 independent experiments.

(E) Bacterial burden of NS398 treated larvae at 4dpi after co-treatment with DMSO or DMOG. Data shown are mean ± SEM, n=33 as accumulated from 3 independent experiments.

