## Supplementary figures and images for "Hif-1α stabilisation polarises macrophages via cyclooxygenase/prostaglandin E2 *in vivo*"

### Figure S1

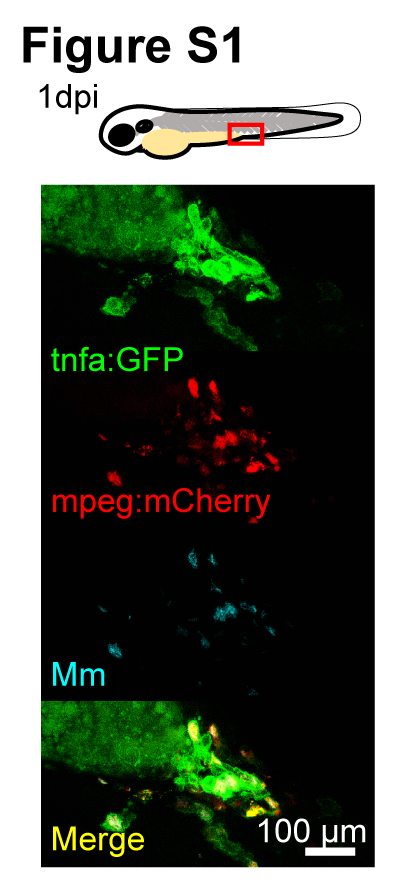

### Figure S2

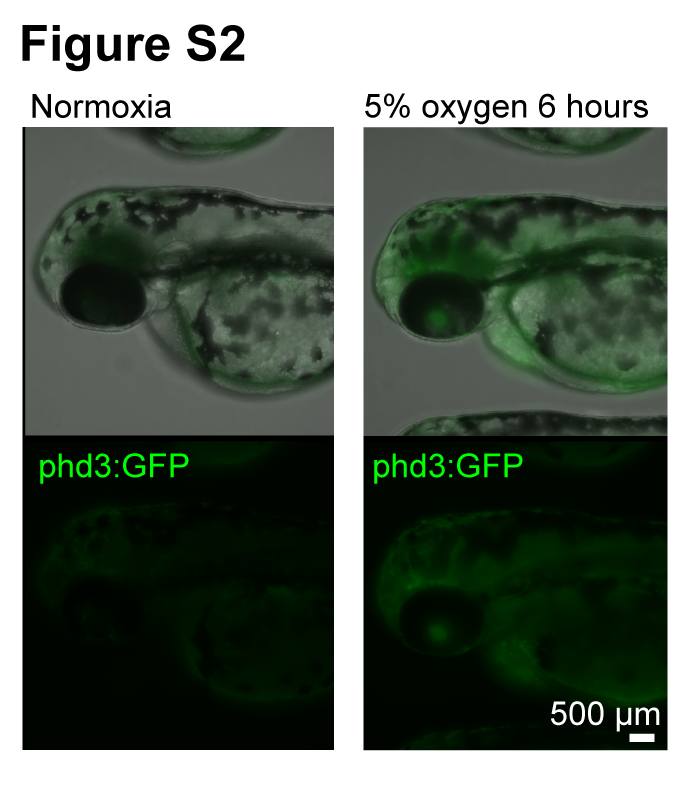

### Figure S3

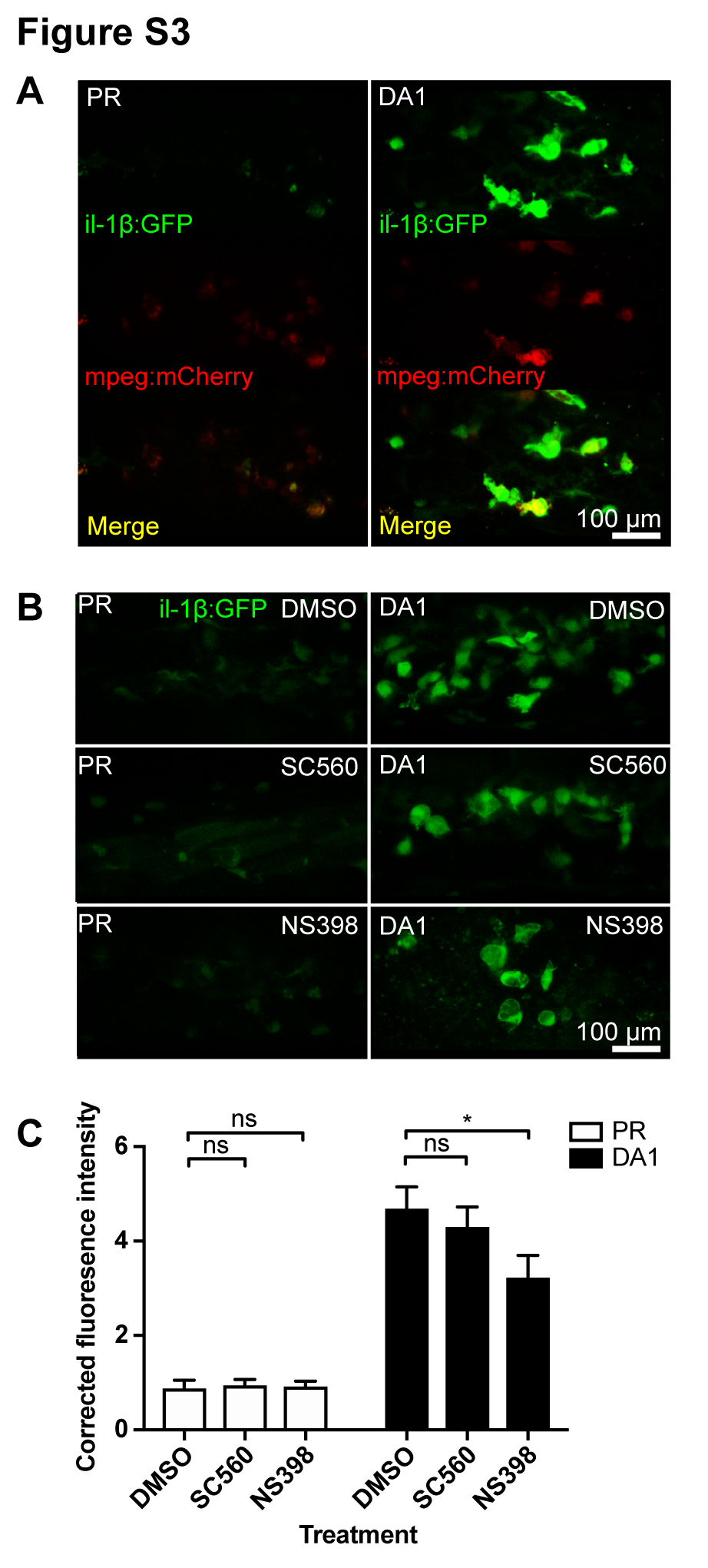

### Figure S4

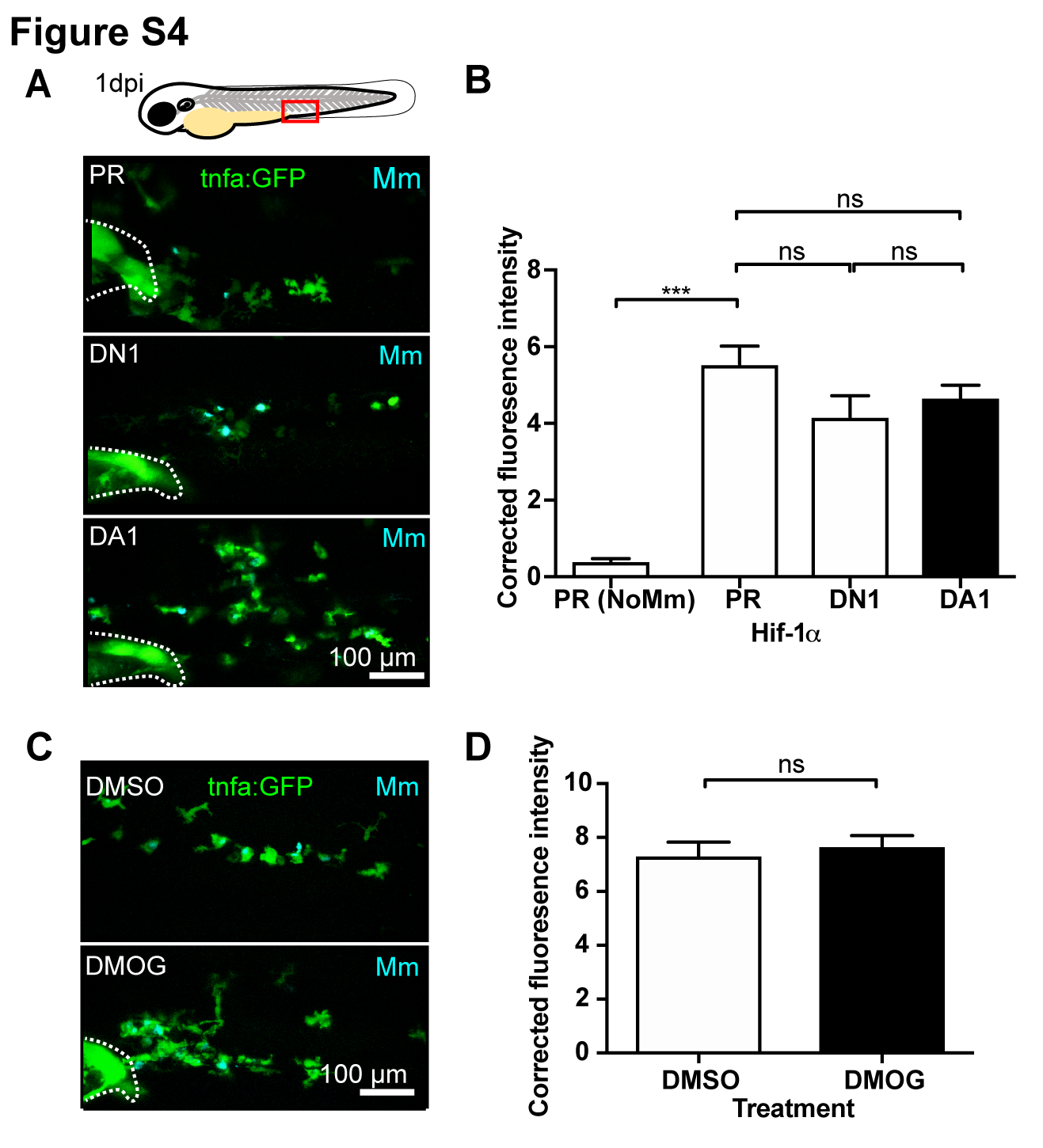

### Figure S5

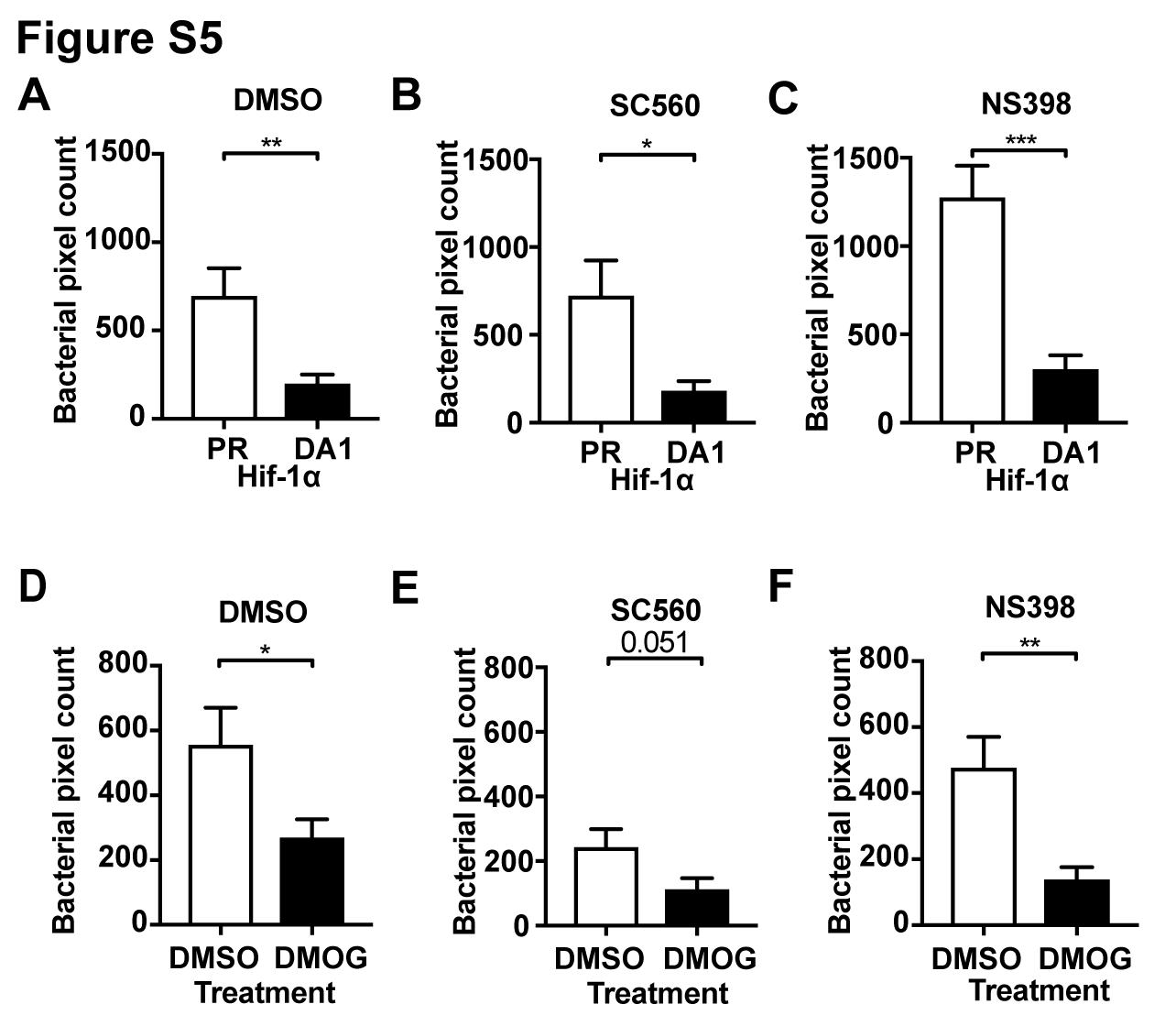
